# Population Analysis of Extracellular Vesicles in Microvolumes of Biofluids

**DOI:** 10.1101/2020.01.10.895037

**Authors:** Joana Maia, Silvia Batista, Nuno Couto, Ana C. Gregório, Cristian Bodo, Julia Elzanowska, Maria Carolina Strano Moraes, Bruno Costa-Silva

**Author notes:** These authors contributed equally to this work (Maria Carolina Strano Moraes, Bruno Costa-Silva).

## Abstract

Extracellular Vesicles (EVs), membrane vesicles released by all cells, are emerging mediators of cell-cell communication. By carrying biomolecules from tissues to biofluids, EVs have attracted attention as non-invasive sources of clinical biomarkers in liquid biopsies. Although frequently employed for content characterization of EVs, the study of bulk preparations lacks information on sub-populations and the intrinsic heterogeneity of vesicles. Importantly, these strategies also difficult the characterization of EVs from small quantities of samples. We here present a Flow Cytometry strategy that enables detailed population analysis of EVs, at the same time decreasing sample volume requirements and accelerating the overall processing time. We show its unique application for quality control of isolates of EVs by comparing the proportion of vesicular and non-vesicular particles in samples prepared by different protocols. In addition, we demonstrate its suitability for the study of populations of EVs from samples characterized by challenging small volumes. To illustrate that, we perform longitudinal non-lethal analysis of EVs in mouse plasma and in single-animal collections of murine vitreous humor. By allowing for the analysis of EVs from minimal amounts of sample, our Flow Cytometry strategy has an unexplored potential in the study of EVs in clinical samples with intrinsically limited volumes. When compared to conventional methods, it also multiplies by several times the number of different analytes that can be studied from a single collection of biofluid.

## Introduction

Extracellular Vesicles (EVs), membrane vesicles released by all cells, are emerging players in cell-cell communication. In addition to differences in size and biogenesis mechanism, EVs are highly heterogeneous in molecular composition and biological properties.^1–4^ Their cargo includes proteins, lipids and nucleic acids, which can be transported both locally and to distant cellular targets through the peripheral circulation. As shown by us^5–8^ and others,^9^ EVs can carry biomolecules from tissues to biofluids and serve as non-invasive sources of liquid biopsies for clinical biomarkers.

The study of EVs often involves the separation of vesicles of similar size (e.g. by ultrafiltration, size exclusion chromatography, nanowire-based traps and deterministic lateral displacement systems), mass and/or density (e.g. by gradient-coupled ultracentrifugation and acoustic separation systems). Once isolated, pooled EVs can be studied for their protein (e.g. by western blotting and mass spectrometry), lipid (e.g. by mass spectrometry) and nucleic acid (e.g. by PCR, DNA and RNA sequencing) content.^10–12^ Despite enabling the characterization of a group of vesicles, these methods lack information on subpopulations and molecular heterogeneity of EVs at a single-vesicle level.

Technologies based on immunoaffinity capture surfaces and beads, such as ELISA and conventional flow cytometry, have enabled the analysis of populations of EVs expressing specific antigens.^2, 11^ However, by capturing heterogeneous groups of EVs containing a common molecule of interest or due to resolution limitations, these methods do not allow for single-vesicle analysis of small EVs.^13–15^ Nanoparticle Tracking Analysis, such as Nanosight, has been used to perform studies of single EVs. However, current protocols cannot study simultaneously single EVs co-stained with more than one dye/fluorescent-conjugated antibody, which limits the study of multiple parameters at a single EV level^16^. Other techniques, such as transmission electron microscopy, perform highly detailed analysis of the structure and antigen expression of single EVs. However, they frequently limit the number of EVs analyzed per sample, complicating the study of large number of EVs and, consequently, the accurate estimation of EVs populations of interest.

By tackling some of these issues, Flow Cytometry (FC) has emerged as a promising tool for thorough single-EV studies. However, existing protocols require EVs’ isolation^1, 17–20^, volume concentration^21^ or immunocapture of subsets of EVs^22^ prior to staining. All these factors increase the processing time, limit the analysis of EVs to sub-populations containing biomolecules of interest and/or prevent sample analysis of small volumes and/or low numbers of EVs.

We here present a strategy that, by not requiring isolation of EVs or concentration prior to staining (Fig.1a), enables analysis of single EVs in both isolated and non-purified samples. We also show its application to quantitatively assess the purity of isolates of EVs prepared by different protocols. Furthermore, we demonstrate its ability to analyze populations of EVs in microvolumes of biofluids, by using it for a longitudinal population study of individual mouse plasma and of individual mouse vitreous humor collections.

## Methods

### Cells

The C57Bl/6 murine PAN02 cell line (also identified as Panc 02, originally induced by 3-MCA^23^) was purchased from the DTP, DCTD Tumor Repository, NIH, and cultured in RPMI 1640 (Corning 15363561,NY, US). The medium was supplemented with 10 % Fetal Bovine Serum (FBS, Biowest S181BH-500, Nuaillé, France) and 1 % penicillin–streptomycin (Gibco 15-140-122, US), and maintained in a humid incubator with 5 % CO_2_ at 37^°^ C. For conditioning, cells were cultured in DMEM, High Glucose, No Glutamine, No Phenol Red (Gibco 31-053-028, US) supplemented with 1 % penicillin–streptomycin and 10 % EV-depleted FBS. FBS was depleted of bovine EVs by ultracentrifugation at 100,000 g for 140 minutes. For the preparation of conditioned medium, 1×10^6^ PAN02 cells were seeded per 150 mm culture dish containing 20 mL of medium, and the conditioned medium was collected after 72 hours of culture.

### Tumor induction, and plasma and vitreous humor collection

All mouse work was performed in accordance with national animal experimentation guidelines (DGAV), animal protocol 0421/000/000/2018. Adult C57Bl/6 female mice (five to eight weeks old) were used for all experimental procedures. Mice were anesthetized using isoflurane 1.5-3 %. Mouse vitreous humor was collected from naïve mice using 29G syringes (BD 324892, New Jersey, US) inserted at a 45° angle into the vitreous cavity 2 mm posterior to the limbus. For tumor induction, a suspension of 1.5×10^6^ PAN02 tumor cells, resuspended in 30 µL Matrigel (Corning 354230, NY, US), was injected intrahepatically with 29G syringes. Blood was collected by retro-orbital bleeding via heparinized glass capillary tubes or by submandibular bleeding via blood collection lancets.

### Purification and characterization of EVs

Mouse blood (3.5 mL) and supernatant fraction of conditioned medium (80 mL) were centrifuged at 500 g for 10 minutes. The collected supernatant was then centrifuged at 3,000 g for 20 minutes (these samples are from now on referred to as “non-purified” – NP – samples), followed by another centrifugation at 12,000 g for 20 minutes. After these initial steps, purification of EVs was performed in plasma by UC-I (bellow) and in conditioned medium according to one of the following protocols (Fig.1a):-(Exoquick) ExoQuick^®^ commercial kit (System Biosciences EXOTC10A-1, Palo Alto, CA), following manufacturer’s instructions;

- (UC-I) 100,000g for 140 minutes, followed by pellet resuspension in 14.5 mL of filtered Phosphate-Buffered Saline (PBS, Corning 15313581, NY, US). This sample was pipetted on top of a 4 mL sucrose cushion (D_2_O containing 1.2 g of protease-free sucrose and 96 mg of Tris base adjusted to pH 7.4), and centrifuged at 100,000 g for 70 minutes. The fraction of interest (4 mL) was aspirated with a 18G needle and taken to a final volume of 20 mL with filtered PBS. The sample was then centrifuged at 100,000 g for 18 hours. The EV-containing pellet was resuspended in filtered PBS;
- (UC-II) 100,000 g for 140 minutes. Pellet was washed with filtered PBS and centrifuged at 100,000 g for 18 hours. The EV-containing pellet was resuspended in filtered PBS;
- (UC-III) 100,000 g for 70 minutes, followed by resuspension of the EV-containing pellet in filtered PBS;
- (UC-IV) 100,000 g for 70 minutes. Pellet was washed with PBS and again centrifuged at 100,000 g for another 70 minutes. The EV-containing pellet was resuspended in filtered PBS.

All solutions used (PBS and sucrose cushion) were sterile (0.22 µm membrane-filtered). The ultracentrifugation (UC) steps were performed in refrigerated conditions (4^°^C) with rotors 50.4Ti or 70Ti (Beckman-Coulter, California, US).

### EVs’ Flow Cytometry

Supernatant fraction of conditioned medium (100 µL), mouse plasma (1 µL in 50 µL of 5,000 U-I-/mL heparin) and vitreous humor (1 µL diluted in 20 µL of 5,000 U-I-/mL heparin) were centrifuged at 500 g for 10 minutes. The collected supernatant was then centrifuged at 3,000 g for 20 minutes. As a simple approach to normalize the input for our staining protocol, all samples were analyzed for particle concentration and size distribution by the NS300 Nanoparticle Tracking Analysis (NTA) system with red laser (638nm) (NanoSight – Malvern Panalytical, UK). Samples were pre-diluted in filtered PBS to achieve a concentration within the range for optimal NTA analysis. Video acquisitions were performed using a camera level of 16 and a threshold between 5 and 7. Five to nine videos of 30 seconds were captured per sample. Analysis of particle concentration per mL and size distribution were performed with the NTA software v3.4.

For staining, 2×10^9^ particles of non-purified (NP) sample or purified EVs were mixed with 40 µL of PBS containing 0.4 µg of anti-CD9 conjugated to phycoerythin (PE) (Thermo Fisher Scientific LABC 12-0091-81, Massachusetts, US) and incubated for 1 hour at 37° C. Samples were incubated with Carboxyfluorescein Diacetate Succinimidyl Ester (CFSE – Thermo Fisher Scientific LTI C34554, MA, USA) in a final concentration of 25.6 µM for 90 minutes at 37° C. The staining with CFSE was done before the addition of the antibody to ensure that the dye would not interfere with the antibody staining. For removal of unbound CFSE and antibody, Size Exclusion Chromatography (SEC) columns (iZON qEV original columns SP1, UK) were used. Samples containing unstained or stained EVs, and appropriate controls, were diluted up to 500 µL of filtered PBS and processed by qEV following manufacturer’s instructions. EVs-enriched fractions #7, #8 and #9 were then pooled and retrieved for analysis with the Flow Cytometer Apogee A60-Micro-Plus (Apogee Flow Systems, UK).

The A60-Micro-Plus machine is equipped with three spatially separated lasers (488 nm – Position C, 405 nm – Position A and 638 nm – Position B), 7 fluorescence color detectors (525/50, LWP590, 530/30, 574/26, 590/40, 695/40, 676/36) and 3 light scatter detectors (SALS, MALS and LALS). More details are available in Table S1.

As internal controls across assays, before each FC experiment we used two commercially available mixes of beads (Apogee 1493 and Apogee 1517, Apogee Flow Systems, UK). Before loading, samples were diluted in filtered PBS to bring their concentration within the operational range of the equipment (maximum of 3,000 events/second). All samples were run at a flow rate of 1.5 µL/minute using a 405 nm – LALS threshold of 70. The 405 nm – LALS PMT was monitored and always maintained below 0.35. This number is an indicator of noise, since it displays the amount of background current, which is a function of the amount of background light reaching the photomultiplier.

For the control experiments (**Figs. S1c, S2a, S2c and S7**), equivalent running times was the stopping criteria utilized. Thus, in these cases control samples were captured for equivalent amounts of time in order to fairly compare the number of positive counts between the different conditions. For the population analysis experiments depicted, the stopping criteria utilized was the number of events acquired, so samples were acquired until a minimum of 250,000 events was reached. In all experimental settings, the data was not pre-gated based on the scatter signals. The acquired data was exported and analyzed with FlowJo software v10.4.2 (FlowJo LLC, US). All flow cytometry files are available at “flowrepository.org” under the Repository ID FR-FCM-Z2EH.

### Calculation of MESF values

The molecules of equivalent soluble fluorochrome (MESF) were calculated for PE and CFSE using SPERO ^TM^ Rainbow Beads Calibration Particles (Spherotech RCP-05-5, USA) according to instructions provided by the manufacturer. In summary, the MFI of the beads was measured with the same acquisition settings applied for the EVs samples. The MFI was converted to relative channel number using the formula (Relative Channel# (#CH) = (R/4)log_10_(MFI), where R is the resolution). Then, the #CH values of the Rainbow beads were plotted against log MESF values and a linear regression was calculated. The MESF values for unknown samples was calculated using the equation corresponding to this linear regression.

### NP-40 Treatment

EVs were lysed by incubation with 2 % NP-40 (Thermo Fisher Scientific 85124, MA, USA) for 1 hour at room temperature, and then stained with CFSE and analyzed by our FC strategy, as described above.

### Statistical analysis

Error bars in graphical data represent means ± s.e.m. Statistical significance was determined using either a two-tailed Student’s t-test or an ANOVA test. A P value under 0.05 was considered statistically significant. Statistical analyses were performed using the GraphPad Prism software (GraphPad Version 7, CA, USA). No statistical method was used to predetermine sample size. The experiments were not randomized, and the investigators were not blinded to allocation during experiments and outcome assessment.

## Results

### Assessment of Flow Cytometry Parameters

Silica and Polystyrene beads were used as reference particles for our EV measurements. Using our settings, it is possible to detect particles that scatter light similarly to 100 nm silica (SiO_2_) beads (**Fig.S1a-i,ii and iii**). In this ApogeeMix, SiO_2_ beads have a refractive index (□) (□ =1.42-1.43) close to that of EVs (□ ~1.39).^24, 25^ Particle concentration and size distribution were verified by Nanoparticle Tracking Analysis (**Fig.S1a-iv**). To identify event coincidence and swarming regime^26^ in our experimental settings, serial dilutions of isolated EVs were performed. The working range was set in the linear region within the operational range indicated by the Apogee A60-Micro-Plus manufacturer (maximum of 3,000 events/second) (**Fig.S1b**).

### Validation of the EVs staining protocol

Our FC strategy relies on the staining of vesicular particles with CFSE (Fig.1a), as previously described in the literature.^17, 19, 27^ A subset of the experiments showed here was performed with conditioned medium, in which cells were grown in medium containing EVs-depleted FBS. To control for the presence of serum-derived vesicles in our samples, serum-free medium and medium containing 10 % of EVs-depleted FBS were stained with CFSE and analyzed using our standard settings, with flow cytometry data being acquired for similar periods of time. CFSE^+^ events count in medium containing 10 % of EVs-depleted FBS were as low as those found in serum-free medium (**Fig. S1c**). For all subsequent analyses, quadrant thresholds were established with unstained and single-stained EVs. Vesicle-free controls containing CFSE, anti-CD9, and both CFSE and anti-CD9 are also shown. Samples were captured using equal time periods, similar to the acquisition times of corresponding samples containing EVs (**Fig.S2a**). To ensure that Anti-CD9 staining was performed in optimal experimental conditions, incremental concentrations of Anti-CD9 were tested (**Fig.S2b**). The working amount of Anti-CD9, was set at 0.4 µg. To certify that the observed CFSE^+^ events were indeed vesicles, isolated EVs were pre-incubated with 2 % NP-40 for 1 hour at room temperature and analyzed using our FC strategy. A reduction of 90 % of CFSE^+^ events was found upon detergent treatment, confirming that most of the observed CFSE^+^ events were EVs (**Fig. S2c**).

Assessment of standardized unit molecules of equivalent soluble fluorochrome (MESF) values allows for cross comparisons between different instruments and laboratories.^28, 29^ In this work, MESF for PE and FITC were then used to determine the fluorescence intensity. The MESF values were measured, with the same acquisition settings applied for all the assays and using a set of Rainbow Beads containing 4 bead populations with known equivalents of FITC molecules and 4 bead populations with known equivalents of PE molecules (**Fig. S3a**). After data collection, the Median Fluorescence intensity (MFI) of these peaks was converted to Relative Channel Number (#CH), and a linear regression of #CH vs log MESF values was performed (**Fig. S3b and c**). This regression allowed for the calculation of MESF values of our experimental controls. In our experimental setting the gates for CFSE^+^ and CD9^+^ were defined such that the CFSE^−^ and CD9^−^ populations displayed approximately 517 and 371 MESF, respectively. The CFSE^+^ population was approximately 10501 MESF whereas the PE^+^ population (CD9) was approximately 1441 MESF (**Fig. S3d**).

### Quantification of populations of EVs in conditioned medium

The quantification of vesicles in conditioned medium (NP) from native tumor cells by our FC strategy showed that ~40 % of particles were CFSE^+^ (**Figs. 1b and S4a**). On the other hand, samples isolated by sucrose cushion-coupled differential ultracentrifugation (UC-I), considered a high-specificity method, contained ~85 % of CFSE^+^ vesicles. Different EVs isolation protocols by ultracentrifugation may utilize different combinations of washing and density flotation steps. Therefore, the impact of these variations in the proportion of CFSE^+^ vesicles in the final isolate was tested (Fig. 1a). With the exception of samples ultracentrifuged overnight without a prior density flotation step (UC-II), in which the proportion of vesicular structures was ~70 %, all other ultracentrifugation-derived preparations contained ~85 % of vesicular structures. As in ultracentrifugation-based methods, ExoQuick^®^ generated samples containing ~80 % of vesicular structures (Fig. 1b).

**Figure 1.**
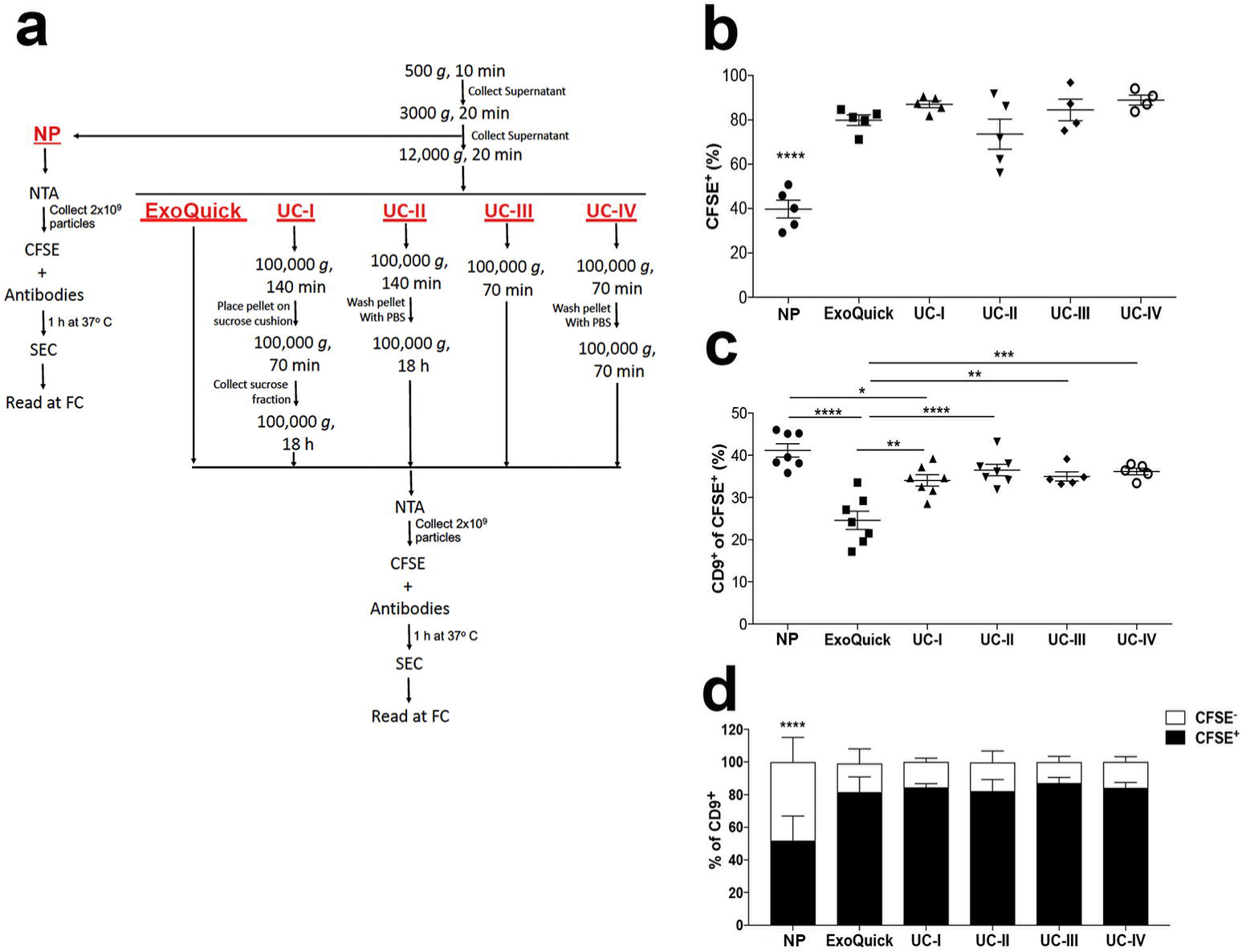
Comparison of isolation methods of EVs. **a**, Flow chart for sample processing by ultracentrifugation (UC), ExoQuick and for analysis of non-purified (NP) samples. **b**, Proportion of CFSE^+^ EVs in non-purified conditioned medium (NP) vs. conditioned medium purified by: ExoQuick^®^; conditioned medium purified by ExoQuick and four distinct protocols ultracentrifugation (UC-I-IV), as indicated in the Figure 1a.****P<0.0001 by ANOVA, with Tukey’s post-test. **c**, Proportion of CD9^+^ events within CFSE^+^ EVs in NP, and medium purified by ExoQuick and ultracentrifugation (UC-I-IV). ****P<0.0001, ***P<0.001, **P<0.01, *P<0.05 by ANOVA, with Tukey’s post-test. **d**, Proportion of CFSE^+^ (black) and CFSE^−^ (white) events within CD9^+^ events in NP samples and samples purified by ExoQuick and ultracentrifugation (UC-I-IV). ****P<0.0001 by ANOVA, with Tukey’s post-test. All data are represented as mean ± s.e.m.

The impact of isolation methods in populations of EVs was also tested. Based on previous reports indicating expression of CD9 in EVs of endosomal origin^4^, CFSE^+^ populations containing this tetraspanin were measured. Although displaying a slight reduction in UC-I, the proportion of CD9^+^ in CFSE^+^ vesicles was comparable between non-purified samples (NP) and those obtained after isolation by ultracentrifugation. On the other hand, samples processed by ExoQuick^®^ contained lower levels of CD9^+^ within CFSE^+^ vesicles (**Figs. 1c and S4b**).

For all processing strategies, detectable levels of CFSE^−^ events displaying CD9 expression were found. We firstly assessed whether these events were due to unbound antibody aggregation or CFSE scarcity. Increments in the concentration of CFSE didn’t increase the proportion of CFSE^+^ EVs (**Fig. S5a**), suggesting that CFSE^−^ vesicles are not the result of sub-optimal experimental conditions for CFSE staining. To exclude the potential presence of aggregates of unbound antibodies, a PBS solution containing anti-CD9 antibody in the concentration used in the staining reaction and pre-cleared by SEC was prepared. Increasing acquisition times were captured to generate a calibration curve. For the acquisition times used in the analysis of conditioned medium-derived EVs, less than 8 % of CFSE^−^CD9^+^ events could be attributed to unbound anti-CD9 (**Fig. S5b and c**). The proportion of CFSE^+^ and CFSE^−^ particles within CD9^+^ particles in samples prepared by different isolation methods was also compared. The number of CFSE^−^CD9^+^ particles was reduced after EVs purification by ExoQuick^®^ or ultracentrifugation (**Figs. 1d and Fig. S5d**), suggesting that the purification methods tested here reduce the presence of CFSE^−^CD9^+^ entities.

### Quantification of populations of EVs in plasma

The proportion of CFSE^+^ vesicles in isolated and non-purified plasma was measured. While ~90 % of particles in isolated samples (UC-I) were CFSE^+^ vesicles, this percentage decreased to approximately 20 % in non-purified samples (NP) (**Figs. 2a and S6a**). Next, the population distribution of CD9 in EVs from mouse plasma was analyzed. Between 15-32 % of CFSE^+^ plasma EVs were CD9^+^. Importantly, this level was comparable between samples that had been isolated by UC-I and in native plasma samples (Fig. 2b). The CFSE^−^CD9^+^ population in these two distinct types of plasma samples was also investigated. In non-purified samples, ~50 % of CD9^+^ events were CFSE^−^. Notably, this proportion decreased to ~30 % after EVs purification by UC-I (**Figs. 2c and S6b**). Thus, according to our estimate of false-positive events (**Fig. S5b**), less than 10% of CD9^+^CFSE^−^ particles can be attributed to unbound anti-CD9 antibodies (**Fig. S6c**).

**Figure 2.**
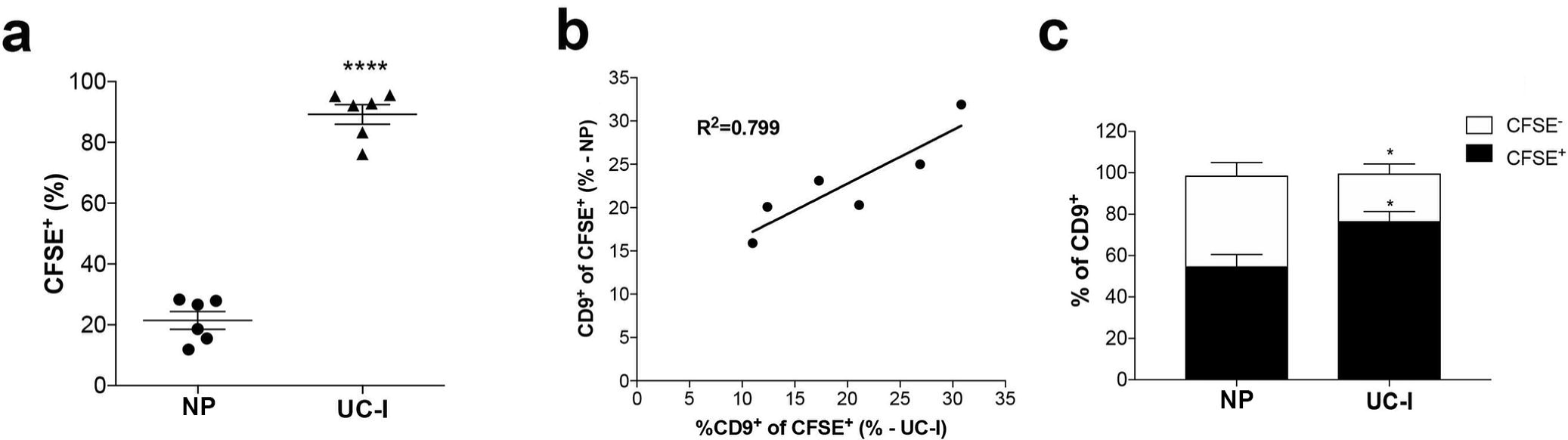
Characterization of EVs in plasma. **a**, Proportion of CFSE^+^ EVs in non-purified plasma (NP) vs. plasma purified by sucrose cushion-coupled ultracentrifugation (UC - I). **b**, Correlation analysis of CD9^+^ events within CFSE^+^ EVs in samples before (NP) and after purification (UC-I). **c**, Proportion of CFSE^+^ (black) and CFSE^−^ (white) within CD9^+^ particles. ****P<0.0001, *P<0.05 differences vs. NP by two-tailed t-test. All data are represented as mean ± s.e.m.

### Longitudinal population studies of EVs in microvolumes of plasma

We tested the applicability of our FC strategy to longitudinal measurements of populations of EVs in microvolumes of non-purified mouse plasma. The presence of tumor cells is expected to modify the plasmatic levels of EVs during the course of the disease. Therefore, plasma from tumor-bearing mice was collected at different time points.^30^ Specifically, blood was collected from each mouse prior to intrahepatic injection of PAN02 pancreatic cancer cells (Day 0), and one (Day 7) and two weeks (Day 14) post-injection, as depicted in Fig. 3a. In this longitudinal analysis of plasma from 10 animals, the proportion of CFSE^+^ events significantly increased in eight of the experimental subjects between days 0-7, in five between days 7-14 and in nine between days 0-14 (**Figs.3b and S7a**). When analyzed in relation to the total concentration of particles present in each time point measured by Nanosight (Fig. 3c), the concentration of CFSE^+^ events *per* µL increased in 6 animals between days 0-7 and days 7-14, and in all animals between days 0-14 (Fig. 3d).

**Figure 3.**
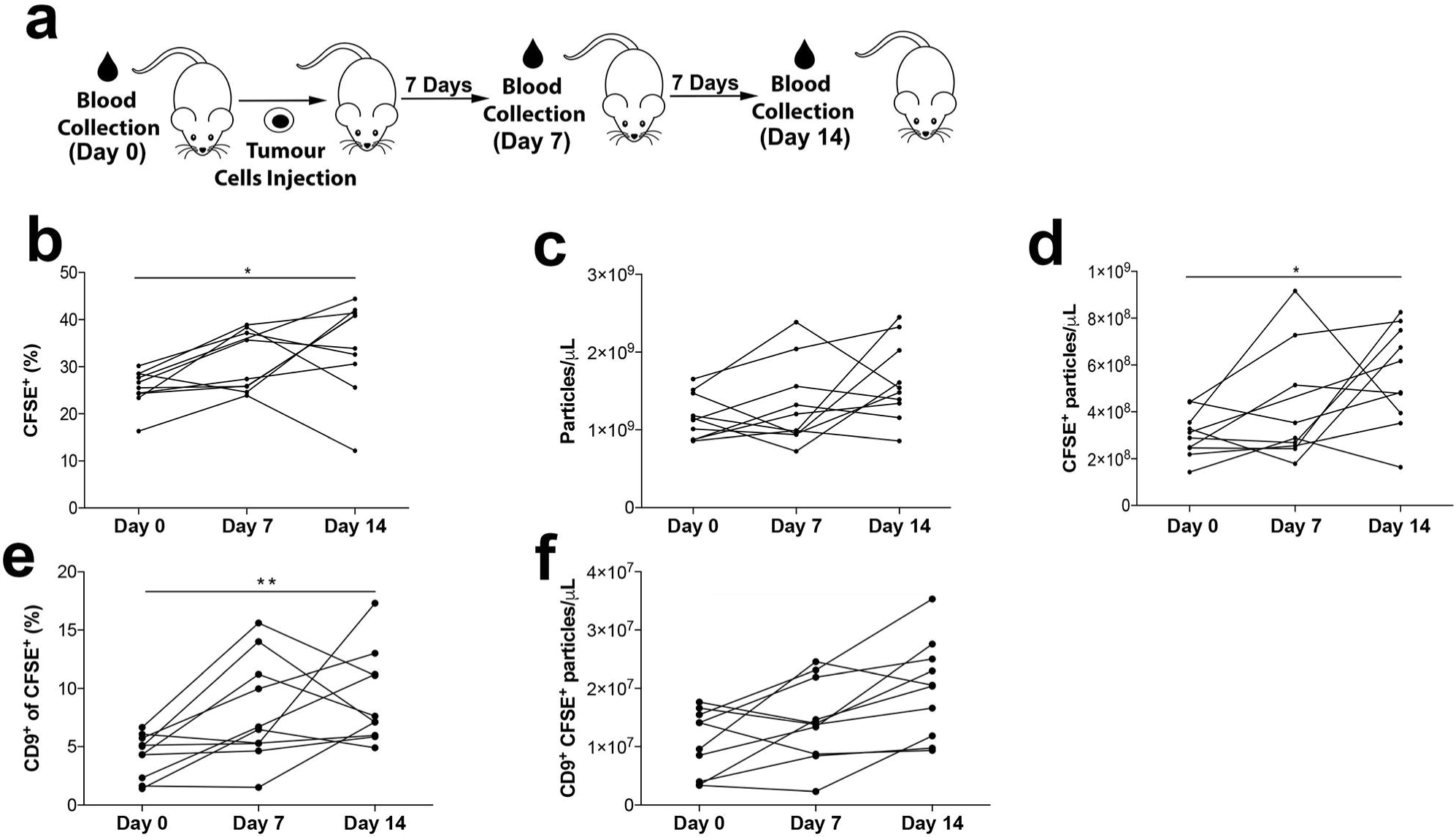
Longitudinal analysis of EVs in plasma. **a**, Experimental setup. **b**, Proportion of CFSE^+^ EVs in non-purified plasma of mice prior (Day 0), and 7 and 14 days after intrahepatic injection of PAN02 tumors. **c**, Concentration of total particles per microliter of plasma, by NanoSight. **d**, Concentration of CFSE^+^ EVs per microliter of plasma. **e**, Proportion of CD9^+^ events within CFSE^+^ EVs. **f**, Concentration of CD9^+^ CFSE^+^ EVs per microliter of plasma. **P<0.01, *P<0.05, differences vs. Day 0 by ANOVA, with Kruskal-Wallis post-test. All data are represented as mean ± s.e.m.

CD9^+^ populations were measured using the experimental settings already described. The proportion of CD9^+^ within CFSE^+^ events increased in 8 animals between days 0-7, in 7 animals between days 7-14 and in 9 animals between days 0-14 (**Figs. 3e and S7b**). Similarly, when analyzing this data in relation to the concentration of particles present in plasma (**Fig.3c**), an increase of CFSE^+^CD9^+^ events *per* µL in 6 animals between days 0-7 and in 9 animals between days 7-14 and 0-14 (Fig.3f) was observed.

### Quantification of populations of EVs in vitreous humor

To further illustrate the application of our FC strategy in performing population analysis of EVs in microvolumes of samples, we used it to study EVs in non-purified vitreous humor, of which 2-2.5 μL *per* mouse/*per* eye was collected. Approximately 68 % of particles were CFSE^+^ vesicles (Fig. 4a and b). However, the levels of CD9^+^ events in vitreous humor (<0.1% - Fig. 4b) were as low as those found in control PBS containing CFSE and anti-CD9 during equivalent sample running times (**Fig. S8**).

**Figure 4.**
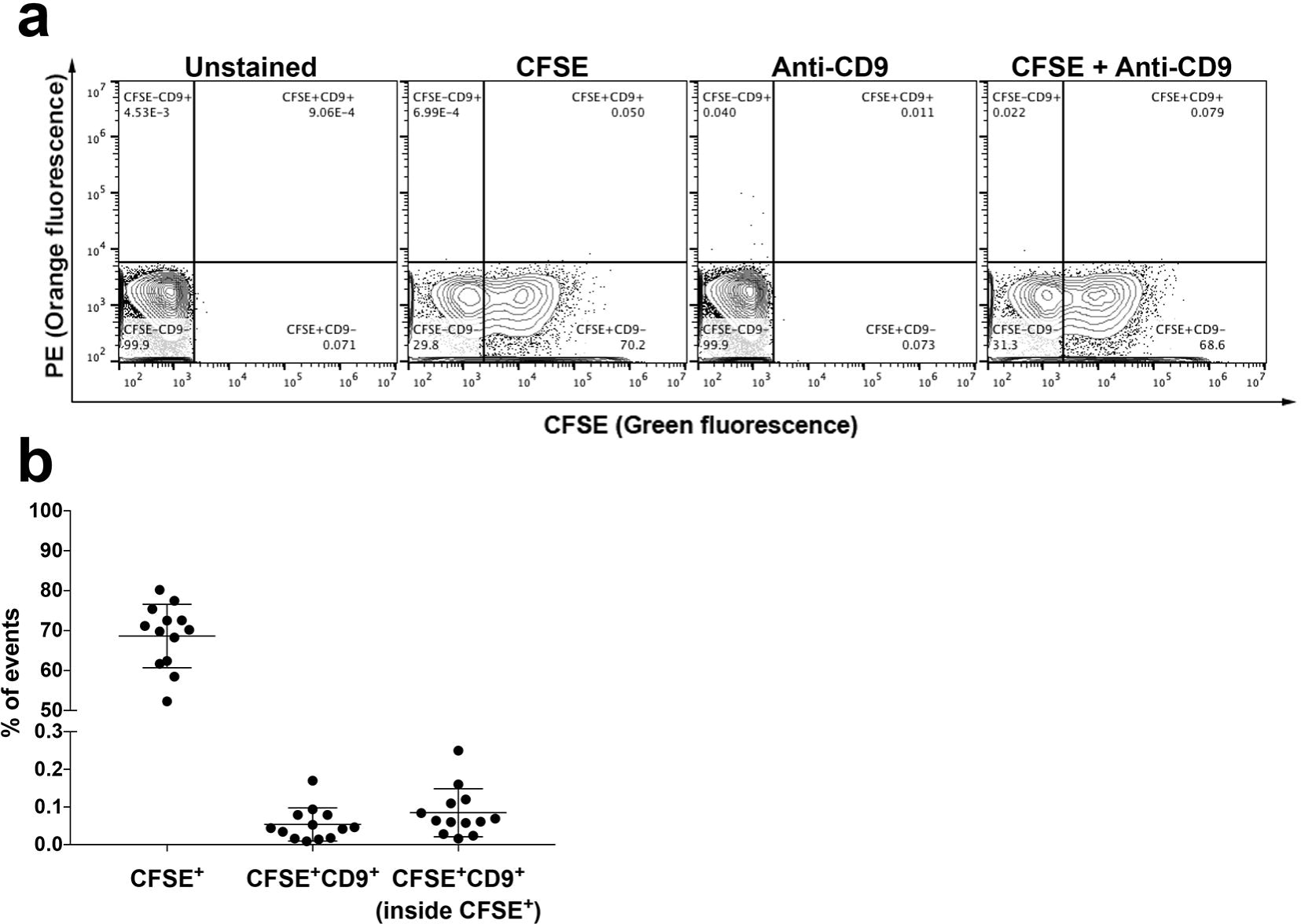
Analysis of EVs in vitreous humor. **a**, Representative plots of unstained and stained EVs from non-purified vitreous humor from naïve mice in single-animal collections, as indicated. **b**, Proportion of CFSE^+^ events, CFSE^+^CD9^+^ events and CD9^+^ events within CFSE^+^ EVs. All data are represented as mean ± s.e.m.

## Discussion

### Measurement of CFSE^+^ events as a strategy to assess sample purity

The lack of consensus regarding methods for purification of EVs remains a challenge, especially when reproducibility between different isolation modalities is desired. This is further complicated by the insufficient means to measure sample purity, notably in complex samples containing a mixture of vesicular and non-vesicular particles, such as plasma. Multiple strategies, such as measurement of protein:particle ratio^31, 32^ and albumin^33, 34^, have been proposed as quality control parameters for the preparation of EVs, particularly when low specificity isolation methods are employed^35^. However, because they fail to provide population information, it is still unclear whether these metrics can accurately reflect the purity of preparations of EVs.

The data here presented supports the vesicular labeling with CFSE as a means for determining quality control of purification protocols of EVs. As shown, the majority of particles present in samples purified by high-specificity methods were CFSE^+^. The employment of ExoQuick^®^ or ultracentrifugation for enrichment of EVs prior to SEC (non-purified) reduced the proportion of CFSE^−^ particles. In addition, lysis of EVs with the detergent NP-40 resulted in a reduction of 90 % of CFSE^+^ events. Together, these results support the usefulness of CFSE as a tool to identify vesicular particles in biofluids, and thus the suitability of our FC strategy for quality control and quantitative comparison of isolates of EVs prepared by different protocols.

This same approach was employed to validate the depletion of EVs from the FBS used in our *in vitro* studies. Previous reports suggest that traces of bovine EVs may persist in the FBS supernatant after depletion steps,^36^ which could interfere in the analysis of EVs from conditioned medium. However, comparison of serum-free medium and medium containing 10 % of EVs-depleted FBS showed an equally reduced proportion of CFSE^+^ events. This indicates that residual FBS-derived EVs was not high enough to impact our analysis. Nonetheless, future studies using our strategy for analysis of EVs in conditioned medium will need to account for background events on a case-by-case basis, especially when using low cell numbers, short conditioning time and/or cells producing low levels of EVs.

### Comparison of purification methods of EVs by FC

In spite of being considered to be a high-recovery and low-specificity method,^35^ isolation based on precipitation polymers such as ExoQuick^®^ resulted in a high proportion of CFSE^+^ EVs. This agrees with studies suggesting that EVs prepared by ultracentrifugation or precipitation polymers are comparable.^37, 38^ In samples prepared by ExoQuick^®^, however, we found that the proportion of CD9^+^ CFSE^+^ events was reduced when compared to non-purified samples and other isolation methods. This suggests that, despite providing a high yield of EVs, ExoQuick^®^ may insert EVs population bias. Precipitating agents have previously been linked to potential loss of biological activity^39^ and structure^40^ of EVs. Thus, our data adds yet another parameter that should be carefully considered before selecting ExoQuick^®^ as a method of choice for the isolation of EVs.

SEC is the technique of choice for many groups interested in studying the composition and biological activity of vesicles, as it allows simple, fast and affordable isolation of EVs. As SEC is a key component of our FC strategy, the proportion of CFSE^+^ particles in our preparations was measured to access the isolation efficacy of EVs by SEC. Although considered a low recovery, high specificity method^35^, conditioned medium processed by SEC contained less than 40 % of CFSE^+^ vesicular particles. This was also the case for more complex samples, such as plasma and vitreous humor, in which the percentage of CFSE^+^ particles after SEC processing were, respectively, ~20 % and ~68 %. These findings agree with recent studies using comparative transmission electron microscopy, in which SEC-derived preparations displayed a lower proportion of structures resembling EVs when compared to samples derived from differential ultracentrifugation.^41^

Differential ultracentrifugation is one of the most commonly used EV purification methods. To improve EV purity, most researchers combine ultracentrifugation with additional techniques following the primary step, such as the use of washing steps with saline and use of density gradients.^42^ We found that the proportion of CFSE^+^ events and CD9^+^ events within the CFSE^+^ gate did not differ in the absence (UC-III) or presence (UC-II and UC-IV) of washing steps with PBS or when a sucrose cushion step was used (UC-I). Our results suggest that these additional steps have no major impact in sample purity. However, a more detailed characterization of the potential impact of these washing and/or separation steps in the selection of populations of EVs with specific composition (protein, sugar, lipid and nucleic acids) will be necessary in future studies.

Together our results show that our FC strategy allows for faster processing times and also substantially decreases the sample volume requirements compared to conventional EVs isolation protocols (Table S2).

### CFSE^−^CD9^+^ events

During the course of this work we observed the presence of CFSE^−^CD9^+^ particles both in conditioned medium and plasma. Although undesirable, protein aggregation may be present in antibody preparations, mainly due to solution conditions such as ionic strength, pH, temperature, pressure and excipients,^43^ and intrinsic properties of antibodies, such as primary sequence, tertiary structure, non-symmetrical hydrophobicity and charge distributions.^44–46^ Therefore, the potential contribution of unbound antibody aggregates to CFSE^−^CD9^+^ events was tested. The vast majority of CFSE^−^CD9^+^ events did not correspond to unbound antibodies and were substantially reduced by isolation of EVs. Results presented by others indicate that CFSE staining could not label 100 % of EVs.^19, 20, 47^ Thus, we reasoned that some of the CFSE^−^ events observed in our experiments could correspond to non-stained EVs. However, unlike the results presented here, in these previous studies CFSE staining was performed in cells before^20^ or during medium conditioning,^19^ and CFSE concentrations were 1.6 to 16 times lower^19, 20, 47^ than those we used. In addition, only ~15 % of the events in samples isolated by differential ultracentrifugation coupled with sucrose cushion (UC-I) were CFSE^−^, and treatment of EVs with the detergent NP-40 caused a reduction of 90 % of CFSE^+^ events, indicating that this dye labels the majority of EVs in our experimental settings. While this may still suggest that CFSE is not capable of staining all EVs present in the sample, it is still unclear to which extent even high-purity isolation methods may provide 100 % pure preparations of EVs. Although our results suggest that the CFSE^−^ events observed in the CD9^+^ population may correspond to non-vesicular particles, future studies, including detailed morphology and composition analysis, will be necessary to further interpret these findings.

### Longitudinal study of EVs in plasma by FC

Longitudinal composition analyses can provide precious temporal information on the dynamics of EVs in physiological and pathological settings.^48–50^ However, these studies are often difficult in microvolumes of samples, mainly due to the limited number of EVs that can be harvested in these experimental conditions.^35^ This constraint frequently leads to insufficient recovery of EVs,^51^ unless small volume samples are pooled from multiple individuals or collections. Moreover, in studies involving small animals, the requirement for lethal bleeding in order to collect enough plasma for the effective isolation of EVs complicates the performance of longitudinal studies and increases the demand for animals, leading to higher costs, higher sample processing complexity and potential bioethical issues. By not requiring isolation of EVs prior to staining, our FC strategy allows for the analysis of both intra- and inter-individual heterogeneity in the population of interest throughout an experiment. In our studies, the proportion of CFSE^+^ EVs and of CD9^+^ events within CFSE^+^ EVs increased in the plasma of mice bearing liver metastatic pancreatic cancer lesions. Based on these results, we are currently studying the potential use of these readouts for follow-up studies of pancreatic cancer patients in the metastatic phase.

### Study of EVs in vitreous humor by FC

The vitreous humor is a small-volume biofluid that contains low protein content, ranging from 120 to 500 ng/µL,^52^ which is frequently considered to arise from filtration of plasma through fenestrated capillaries of the ciliary body stroma via the iris root.^53^ Besides the quantitative differences in protein content, a comparison of vitreous humor and plasma proteome revealed that only 58% of the vitreous humor proteins have also been identified in human plasma.^52^ Consistent with this, our analysis revealed that vitreous fluid contains three times more CFSE^+^ vesicular structures when compared to plasma. Furthermore, it contained insignificant levels of CD9^+^ events, comparable to those found in control solutions with only CFSE and anti-CD9. These results reinforce the idea that the CFSE^−^CD9^+^ particles we observed in conditioned medium and plasma are not an artifact, and are consistent with the previously reported absence of CD9 in EVs from vitreous humor.^54–56^. However, it is still unknown whether the absence of CD9^+^ vesicles is a result of the filtration that occurs during the production of vitreous humor, uptake and degradation of this vesicle population by ocular cells, higher prevalence of non-endosomal EVs and/or other mechanisms. Although it is unclear to which extend ocular cells contribute to the collection of EVs found in vitreous humor, our FC strategy can be potentially used to study these vesicles both in pre-clinical and clinical settings as potential biomarkers and biological mediators of eye diseases.

## Conclusion

By allowing for the analysis of conditioned medium volumes as small as 100 µL, our FC strategy can be potentially used to characterize the heterogeneity of EVs and the differential packaging of biomolecules during the biogenesis of EVs in highly controlled small-scale *in vitro* systems. This approach also makes it possible to study EVs populations of EVs from as little as ~1 µL of plasma or vitreous humor. By doing so, our strategy represents a precious tool to identify novel physiological and pathological cell-to-cell communication systemic networks involving EVs in *in vivo* models. Our FC strategy has an unexplored capability to be applied in the study of populations of EVs in native clinical samples, multiplying the number of different analysis from a single biofluid collection. It also has a great potential to enable the study of populations of EVs in samples with intrinsically limited volumes, such as lacrimal, vitreous humor and synovial fluids, facilitating the use of these biofluids as liquid biopsies on clinical settings.

## Supporting information

MIFlowCyt-EV

## Acknowledgements

We thank A. Vieira and D. Fortunato, for helpful discussions and technical support, and Dr. M. de Sousa for her interest in this work and in the preparation of the manuscript. J.M. was supported by “Fundação para a Ciência e a Tecnologia” (PD/BD/105866/2014). S.B. was supported by the EMBO Installation Grant 3921. A.G. was supported by the grant 2017NovPCC1058 from Breast Cancer Now’s Catalyst Programme, which is supported by funding from Pfizer. J.E. was supported by the grant 765492 from H2020-MSCA-ITN-2017. This work was supported by the Champalimaud Foundation, the grant 751547 from H2020-MSCA-IF-2016 and the grant LCF/PR/HR19/52160014 from “La Caixa” Foundation.

## Author contributions

J.M. designed and performed all experiments. S.B., N.C. and A.G. contributed to animal experimentation. C.B. contributed to sample processing, EVs characterization and equipment maintenance. J.E. contributed with the Pan02 EVs samples for Figure S2a Figure S2c. M.C.S.M. built the experimental setup. B.C.S. conceived the project. All authors wrote and reviewed the manuscript.

---

**Supplementary Figure S1 | EVs’ Flow Cytometry parameters. a**, Flow cytometry analysis of a mix of polystyrene (PS) and silica (Si) beads. Depicted in the histograms are: 110 nm PS, 180 nm Si, 240 nm Si, 300 nm Si (i) and 83 nm PS and 100 nm Si beads (ii) in the context of purified PAN02 EVs (iii). The vesicular particle size distribution, analyzed by NanoSight, is also shown (iv). **b**, Analysis of events/µL in relation to sample dilution factor. The utilized working range is indicated. **c**, Representative plots of CFSE-labeled serum-free and 10% EV-depleted FBS medium; the indicated counts (CFSE+) correspond to the events within the quadrant.

**Supplementary Figure S2 | Internal controls and detergent lysis of EVs. a**, Representative plots of buffer containing CFSE only, anti-CD9 only, and CFSE and anti-CD9 (upper panels), and of unstained EVs, EVs stained with CFSE, EVs stained with anti-CD9, and EVs double stained with CFSE and anti-CD9 (lower panels); the indicated counts correspond to the events within the upper right quadrant (CFSE^+^CD9^+^). **b**, Titration of Anti-CD9 staining. **c**, Detergent lysis of EVs. The plots are representative of unstained and non-lysed EVs, CFSE-stained non-lysed EVs, unstained EVs lysed with NP-40 and CFSE-stained EVs lysed with NP-40; the counts of CFSE^+^ particles are indicated. The graph indicates the reduction in the percentage of CFSE^+^ EVs after lysis with NP-40.

**Supplementary Figure S3 | Assessment of MESF values. a**, Histograms of the RCP-05-5 beads in the FITC and PE channel. **b**, The FITC and PE Median Fluorescence Intensity (MFI) of each peak was measured and converted to Relative Channel Number (#CH), and the MESF values calculated (using template provided by manufacturer). **c**, Calibration graphs and linear regression where the MFI is plotted in the x-axis and log MESF is plotted in the y-axis. **d**, Calculation of FITC and PE MESF values for EVs samples stained with CFSE or Anti-CD9.

**Supplementary Figure S4 | Different purification methods of conditioned medium samples. a**, Representative plots of CFSE-labeled particles from non-purified conditioned medium (NP), and from conditioned medium purified by ExoQuick or by distinct ultracentrifugation protocols (UC-I-IV). **b**, Representative plots of particles labeled with CFSE and anti-CD9 from NP, and from conditioned medium purified by ExoQuick or by ultracentrifugation (UC-I-IV). The lower panels indicate the CD9 status within CFSE^+^ EVs.

**Supplementary Figure S5 | Analysis of false-positive CD9^+^ events in conditioned medium. a**, Titration of CFSE staining. **b**, Calibration curve of false-positive events. Buffer containing anti-CD9, in the same concentration used in the staining reaction and pre-cleared by SEC, was analyzed by Flow Cytometry for increasing acquisition times. **c**, Estimate percentage of false-positive events by unbound anti-CD9 within CFSE^−^CD9^+^ in each purification context. **d**, Representative plots of particles labeled with CFSE and anti-CD9 from NP, and from conditioned medium purified by ExoQuick or by ultracentrifugation (UC-I-IV). The lower panels indicate the CFSE status within CD9^+^ EVs.

**Supplementary Figure S6 | Analysis of CD9^+^ plasma particles. a**, The plots are representative of CFSE-labeled particles from non-purified (NP) plasma, or from plasma purified by ultracentrifugation (UC-I). **b**, Representative plots of particles labeled with CFSE and anti-CD9 from NP and UC-I plasma; the lower panels indicate the CFSE^+^ and CFSE^−^ events within CD9^+^ particles. **c**, Estimate percentage of false-positive events by unbound anti-CD9 in each experimental setting.

**Supplementary Figure S7 | Longitudinal analysis of plasma EVs. a**, Representative plots of CFSE-labeled EVs from non-purified plasma of mice prior (Day 0), and at 7 and 14 days after intrahepatic injection of PAN02 cells. **b**, Representative plots of particles from NP plasma double labeled with CFSE and anti-CD9; the lower panels indicate the CD9^+^ and CD9^−^ particles within CFSE^+^ EVs.

**Supplementary Figure S8 | CD9^+^ events in vitreous humor.** Representative plots of PBS and vitreous humor double labeled with CFSE and anti-CD9. Samples were captured during 250 seconds. The indicated counts (CFSE^+^CD9^+^) correspond to the events within the upper right quadrant.

**Supplementary Table S1: Apogee A60 configuration and laser power**

**Supplementary Table S2: Comparison of Sample Volume and Processing times between isolation methods of EVs**

