## Supplemental Table 1 for "Population Analysis of Extracellular Vesicles in Microvolumes of Biofluids"

**Table S1: Apogee A60 configuration and laser power**

| **Channel Number** | **Short Channel Name** | **Full Channel Name** | **Optical Filter Name** | **Laser Wavelength** | **Laser**  **Power** | **PMT**  **Voltage** |
| --- | --- | --- | --- | --- | --- | --- |
| Ch1 | 405-SALS | Small Angle Light Scatter |  | 405nm | 200mW | 400V |
| Ch2 | 405-LALS | Large Angle Light Scatter |  | 405nm | 200mW | 400V |
| Ch3 | 405-Grn | Green Fluorescence | BP-525/50 | 405nm | 200mW | 500V |
| Ch4 | 405-Org | Orange Fluorescence | LWP-590/35 | 405nm | 200mW | 500V |
| Ch5 | APC | Red Fluorescence | BP-676/36 | 638nm | 150mW | 550V |
| Ch6 | CFSE | Green Fluorescence | BP-525/50 | 488nm | 200mW | 525V |
| Ch7 | PE | Orange Fluorescence | BP-575/30 | 488nm | 200mW | 500V |
| Ch8 | 488-Red | Red Fluorescence | BP-676/36 | 488nm | 200mW | 500V |
| Ch9 | 488-DRed | Deep Red Fluorescence | LWP-740 | 488nm | 200mW | 500V |
