## Supplementary material for "Population Analysis of Extracellular Vesicles in Microvolumes of Biofluids": MIFlowCyt-EV

**MIFlowCyt -EV Framework**

**1. Preanalytical variables & Experimental design**

**1.1 Preanalytical variables conforming to MISEV guidelines.**

The sources of the EVs used in this work were Pan02 Cell Line conditioned medium, plasma and vitreous humor from adult C57Bl/6 female mice (five to eight weeks old). The Pan02 Cell Line conditioned medium was collected after 72hours of conditioning (see conditioning procedure in Supporting Information, Experimental Details, Cells). Mouse Blood was collected by retro-orbital bleeding or by submandibular bleeding and the Mouse vitreous humor was collected using 29G syringes inserted into the vitreous cavity. Supernatant fraction of conditioned medium, mouse plasma and vitreous humor were centrifuged at 500 g for 10 minutes. The collected supernatant was then centrifuged at 3,000 g for 20 minutes. After these initial steps, purification of EVs was performed in plasma and in conditioned media according the protocols in Supporting Information, Experimental Details, Purification and characterization of EVs. All samples were stored at -80^o^C.

**1.2 Experimental design according to MIFlowCyt guidelines**.

The aim of this work was to present a strategy that by not requiring isolation of EVs or concentration prior to staining, enables population analysis of EVs in biofluids with unpreceded small volumes. The samples were analyzed on an Apogee A60-Micro-Plus at a flow rate of 1.5 µL/minute. The variables in our FC experiments were: EVs staining with CFSE, EVs staining with CD9, EVs samples prepared with different protocols, different sample volume requirements.

**2. Sample Preparation**

**2.1 Sample staining details**

For our FC strategy staining, 2x10^9^ particles of unprocessed sample or purified EVs were mixed with 40 µL of PBS containing 0.4 µg of anti-CD9 conjugated to phycoerythin (PE) (Thermo Fisher Scientific - LABC 12-0091-81, Massachusetts, US) and incubated for 1 hour at 37° C. Next, samples were incubated with Carboxyfluorescein Diacetate Succinimidyl Ester (CFSE – Thermo Fisher Scientific - LTI C34554) (6µL of 196µM CFSE working solution) in a final concentration of 25.6 µM and incubated for 90 minutes at 37° C. For removal of unbound CFSE and antibody, Size Exclusion Chromatography (SEC) columns (iZON - qEV original columns SP1, UK) were used. Samples containing unstained or stained EVs, and appropriate controls, were diluted up to 500 µL of filtered PBS and processed by qEV following manufacturer’s instructions. EVs-enriched fractions #7, #8 and #9 were then compiled and retrieved for analysis with the Flow Cytometer Apogee A60-Micro-Plus (Apogee Flow Systems, UK).

**2.3 Sample dilution details**

Before being analyzed in the Apogee A60-Micro-Plus, samples were diluted in filtered PBS (0.22 µm membrane filters) to bring their concentration within the operational range of the equipment (maximum of 3,000 events/second).

**3. Assay Controls**

**3.1 Buffer alone controls**

In our experimental settings, we measured filtered PBS with the same flow cytometer and acquisition settings as the stained samples. The mean count rate was approximately 10-30 events per second, which is substantially lower than the mean count rates (~1000 events per second) for stained samples. When any this data of the Buffer-only controls was utilized, these samples were recorded for the same period of time as the samples they were compared to (see Supporting Information, FigS2a).

**3.2 Buffer with reagent controls**

In our experimental settings, we analyzed filtered PBS with CFSE, filtered PBS with Anti-CD9 and PBS with CFSE and Anti-CD9, which were measured with the same flow cytometer and acquisition settings as the stained samples (see Supporting Information, FigS2a.)

**3.3 Unstained controls**

In all of our experimental settings Unstained samples were measured with the same flow cytometer and acquisition settings as the stained samples (see Supporting Information, FigS2a).

**3.5 Single-stained controls**

In all of our experimental settings Single-Stained samples (stained only with CFSE or stained only with Anti-CD9) were measured with the same flow cytometer and acquisition settings as the stained samples (see Fig4a and Supporting Information, FigS2a). For instance, in Supporting Information, FigS2a, we obtained on approximately 517 MESF for CFSE^-^ and 10501 MESF for CFSE^+^, and approximately 371 CD9^-^ MESF and 1441 CD9^+^ MESF.

**3.7 Serial Dilution**

To identify event coincidence and swarming regime in our experimental settings, serial dilutions of isolated EVs were performed. The working range was set in the linear region: maximum of 3,000 events/second (see Supporting Information Fig.S1b).

**4. Instrument Calibration and Data acquisition**

**4.1 Trigger Channel(s) and Threshold(s)**

All samples were run at a flow rate of 1.5 µL/minute using a 405nm – LALS threshold of 70. The 405 nm – LALS PMT was monitored and maintained bellow 0.35. The flow cytometer configuration, laser powers and PMT voltage are available in Supporting Information, Table S2.

**4.2 Flow Rate / Volumetric quantification**

The samples were analyzed on an Apogee A60-Micro-Plus at a flow rate of 1.5 µL/minute. Because the A60-Micro is equipped with a syringe pump with volumetric control, we selected a flow rate of 1.5 µL/minute for all measurements, as suggested by the manufacturer.

**4.3 Fluorescence Calibration.**

The molecules of equivalent soluble fluorochrome (MESF) were calculated for PE and CFSE, using SPERO TM Rainbow Beads Calibration Particles (RCP-05-5, Sperotech, USA) and according to the template provided by the manufacture. In summary, the MFI of the beads was measured in the same acquisition settings applied for the EVs samples. The MFI was converted to relative channel number using the formula (Relative Channel# (#CH) = (R/4)log10(MFI), where R is the resolution). Then, the #CH values of the Rainbow beads were plotted against log MESF values and a linear regression was calculated. The calibration curves and linear regressions are shown in Supporting Information, FigS3b and FigS3c. In our experimental setting, the CFSE^-^ population displayed approximately 517 MESF and CD9^-^ population approximately 371 MESF.

**6. FC Data Reporting**

**6.1 Completion of MIFlowCyt checklist.**

The MIFlowCyt checklist is available bellow in TableM1.

**Table M1. MIFlowCyt checklist**

| **Requirement** | **Please Include Requested Information** |
| --- | --- |
| 1.1 Purpose | Present a strategy that by not requiring isolation of EVs or concentration prior to staining, enables population analysis of EVs in biofluids with unpreceded small volumes. |
| 1.2 Keywords | microvolume, extracellular vesicles, flow cytometry, population study, sample purity |
| 1.3 Experimental variables | EV staining with CFSE, EV staining with CD9, EVs samples prepared with different protocols, different sample volume requirements |
| 1.4 Organization name and address | Champalimaud Centre for the Unknown Av. Brasília 1400-038 Lisbon, Portugal |
| 1.5 Primary contact name and email address | Bruno Costa-Silva, Maria Carolina Strano Moraes, |
| 1.6 Date and time period of the experiment | July 2017-December 2019 |
| 1.7 Conclusions | Our FC method allows for quality control of isolates of EVs and the study of populations of EVs in samples with small volume available, (e.g. non-lethal longitudinal studies and single-animal collections of mice vitreous humor). By lowering the sample requirement, our FC method multiplies the number of different analytes that can be studied. |
| 1.8 Quality Control measures | The molecules of equivalent soluble fluorochrome (MESF) were calculated for PE and CFSE, using SPERO TM Rainbow Beads Calibration Particles (RCP-05-5, Sperotech, USA) and according to the template provided by the manufacture. In summary, the MFI of the beads was measured in the same acquisition settings applied for the EVs samples. The MFI was converted to relative channel number using the formula (Relative Channel# (#CH) = (R/4)log10(MFI), where R is the resolution). Then, the #CH values of the Rainbow beads were plotted against log MESF values and a linear regression was calculated. Using the resulting equation, it was possible to calculate the MESF values for unknown samples. |
| 1.9 Other relevant experiment information | ---- |
| 2.1.1.1 Sample description | Pan02 Cell Line conditioned medium, plasma and vitreous humor from mouse |
| 2.1.1.2 Biological sample description | Pan02 Cell Line- Murine pancreatic adenocarcinoma cell line Plasma and Vitreous Humor from adult C57Bl/6 female mice (five to eight weeks old) |
| 2.1.1.3 Biological sample source organism description | Plasma and Vitreous Humor from adult C57Bl/6 female mice (five to eight weeks old) |
| 2.1.2.2 Environmental sample location | All samples were stored at -80oC |
| 2.2 Sample Characteristics | Pan02 Cell Line conditioned medium is expected to contain EVs. Platelet free Plasma and Vitreous Humor from mouse is expected to contain EVs, lipoproteins and proteins |
| 2.3 Sample treatment description | Pan02 Cell Line conditioned medium was collected after 72hours of conditioning (see conditioning procedure in Supporting Information, Experimental Details, Cells). Supernatant fraction of conditioned medium, mouse plasma and vitreous humor were centrifuged at 500 g for 10 minutes. The collected supernatant was then centrifuged at 3,000 g for 20 minutes. After these initial steps, purification of EVs was performed in plasma and in conditioned media according the protocols in Supporting Information, Experimental Details, Purification and characterization of EVs.  The staining procedure was done as described in Supporting Information, Experimental Details, EVs Flow Cytometry. |
| 2.4 Fluorescence reagent (s) description | Anti-CD9 conjugated to phycoerythin (PE) (Thermo Fisher Scientific - LABC 12-0091-81, Massachusetts, US); Carboxyfluorescein Diacetate Succinimidyl Ester (CFSE – Thermo Fisher Scientific - LTI C34554) |
| 3.1 Instrument manufacturer | Apogee, Hemel Hempstead, UK |
| 3.2 Instrument model | A60-Micro-Plus |
| 3.3 Instrument configuration and settings | The samples were analyzed at a flow rate of 1.5 µL/minute on an Apogee A60-Micro-Plus, using a 405nm – LALS threshold of 70. The 405 nm – LALS PMT was monitored and maintained bellow 0.35. The A60-Micro-Plus machine was equipped with three spatially separated lasers (488 nm – Position C, 405 nm – Position A and 638 nm – Position B), 7 fluorescence color detectors (525/50, LWP590, 530/30, 574/26, 590/40, 695/40, 676/36) and 3 light scatter detectors (SALS, MALS and LALS). More details are available in Supporting Information in Table S2. For the control experiments (Supporting Information, Figs. S1c, S2a, S2c and S7), equivalent running times was the stopping criteria utilized. Thus, in these cases samples were captured for equal times, in order to fairly compare the number of positive counts between the different conditions. For the population analysis experiments depicted, the stopping criteria utilized was the number of events acquired, so samples were acquired until a minimum of 250,000 events was reached. In all experimental settings, the data was not pre-gated based on the scatter signals. The acquired data was exported and analyzed with FlowJo software v10.4.2 (FlowJo LLC, US). |
| 4.1 List-mode data files | May be requested by emailing |
| 4.2 Compensation description | A compensation matrix was applied after the data collection, since the fluorophore combinations used displayed overlapping emission spectra. The compensation matrix was designed in the FlowJo software v10.4.2, during data analysis. |
| 4.3 Data transformation details | No data transforms were applied |
| 4.4.1 Gate description | The acquired data was exported and analyzed with FlowJo software v10.4.2. Gates were based on fluorescence of a blank sample (unstained), samples labeled only with CFSE and samples labeled only with Anti-CD9 (See Supporting Information, FigS2a). The lower boundaries of the fluorescent gates were determined, resulting in approximately 517 MESF for CFSE- and 371 CD9- MESF. |
| 4.4.2 Gate statistics | Percentages of events within each gate of the contour plot represent percentage of the total population or inside CFSE+ population (which are properly signalized) |
| 4.4.3 Gate boundaries | Images of the gates can be seen in the contour plots within the manuscript and Supporting Information, FigS2a. |
